# Time Evolution of the Stroke Symptom-Herb Networks Based on TCM Prescriptions

**DOI:** 10.1101/2021.08.23.457415

**Authors:** Xing Fu, Qianqian Li, Feng Yang, Ranran Zhou, Ran Li, Jianchen Hou, Xiaohua Tao

**Affiliations:** Department of Shanghan, college of traditional Chinese medicine, Beijing University of Chinese Medicine, Beijing, China

**Keywords:** TCM prescription, Stroke, Stroke Symptom-Herb Networks, Time series analysis

## Abstract

Traditional Chinese Medicine (TCM) has its origins in distant antiquity and has piled up over a long time with much knowledge about diseases, especially stroke. Different combinations of symptom variables yield different combinations of herbs to form a myriad of prescriptions, and they have undergone repeated confirmation and are worthy objects of excavation and analysis. Herbal studies on stroke have developed from genomics to transcriptomics, proteomics and metabolomics, yet more thought is needed on putting time in a wider perspective of the symptoms and herbs for stroke in Chinese medicine.

Due to this, we studied the dynamic structure of TCM prescriptions on stroke, using over 270 TCM prescription books containing 2231 prescriptions related to stroke recorded from 341 to 2000 CE. We labeled the functions of the prescriptions with the symptoms based on subject terms in MESH *Neurologic Manifestations*, then standardized the herbs in the prescriptions, and finally connected the co-occurring symptoms and herbs in the prescriptions to build an undirected complex network. The Stroke Symptom-Herb Networks (SSHNs) can be seen from its network characteristics that it is not a random network and has small-world characteristics. It has experienced two peaks in its nearly 1700-year history, during the Song dynasty, the Ming and Qing dynasties. From 600 years onwards, the core herb cluster has been initially formed. The comparison of sub-network similarities allowed us to identify several symptoms with similar herb clusters.

We divided the community based on modularity, and by analyzing the community evolution, we found a more fixed historical evolutionary trend with *Hemiplegia* and *Sialorrhea* nodes and their associated symptom and herb nodes. In the time series analysis, we found many symptom-herb combinations that were consistently closely related to historic time depends on assessing the similarity between the symptoms and the herbs. The complex network provides a distinctive perspective for understanding the symptom-herb relationships embedded in TCM prescriptions in remote antiquity.

## Introduction

Chinese medicine has gathered over a long time much knowledge about disease, especially stroke. The TCM Ancient Books named *Zhouhou Beiji Prescription*, written in 341 CE, was the first Chinese monograph for clinical first-aid and one of the earliest extant prescription books that clearly describes the TCM patterns of stroke, such as *Hemiplegia*, *Facial Paralysis*, and *Voice Disorders*, etc. Since then prescription books have continuously and plentifully recorded the TCM patterns and herbs of stroke, thus forming a complex and extensive network of symptoms and herbs. At the same time, the network has further evolved and developed in the historical run, forming a time series of stroke symptoms and herbs. Therefore, besides to networks, another key focus of our study is time, from which we believe the rules of developmental change in stroke symptoms and herbs can be identified and revealed.

Network science is the study of network models, and it does not suddenly appear when it was realized in the mid-1990s that networks could be models of complex systems[1]. Behind every complex system, there is an intricate network, and networks portray the interrelationships between the components of complex systems. Network science is dedicated to exploring the universal laws that characterize the widespread networks of nature. Many widely varying networks in nature or society, the nodes of the World Wide Web are web pages, and the links between web pages are Uniform Resource Locators (URLs) used by computer algorithms to resolve addresses; the nodes of social networks are people, and the links include relationships such as family, workplace and friends. A key finding of network science is the networks that form in these different contexts are similar in their architecture, suggesting they form under to guide the same principals. Because of its universality, networking is destined to be a crosscutting science involving mathematics, physics, social science, computer science, and other disciplines[2]-[3]. Network science offers methods and techniques to study the intricate relationship between symptoms and herbs in stroke, analyzing the vast amount of relationships and data in complex networks with their system’s structure and dynamics, and presenting them in a visual way.

Nevertheless, the long history of time has also led to barriers to communication. The main reason TCM patterns are hard to match with modern medical symptoms is that most TCM patterns are recorded in ancient Chinese, and only TCM clinicians who have studied ancient medical literature can achieve a real understanding of TCM patterns. Yet even fewer of them can accurately understand modern medical symptom terminology. This work must be done by clinical experts in cerebrovascular disease who are trained in TCM and familiar with modern stroke terminology.

In this paper, firstly, we screened the stroke and its synonyms to form the stroke prescription database containing 270 prescription books, and standardized the functions and compositions of the stroke prescriptions in the database to form the terminology of TCM patterns and standardized herbal terms. Second, we screened and expanded symptoms according to their definitions based on subject terms within MESH *Neurologic Manifestations*. Then, we invited experts in TCM stroke to match all symptoms with TCM patterns, thus completing the symptom labeling of prescriptions for their functions. Finally, we connected the symptoms and herbs co-occurring under the same prescription, thus building a complex network of stroke symptom-herb. By analyzing their particular variations in time, we discovered some details and patterns that hide in time.

## Materials and Methods

### Terminology standardization and matching

As one can view in Figure 1, we sifted through 270 prescription books for stroke and its synonymous from our TCM ancient books indexing platform (http://114.255.40.130:60080/metaservice2/) containing over 600 ancient Chinese medical books. First, on TCM ancient books indexing platform, the functions and compositions of the prescriptions are separated and labeled manually. Each exported prescription information contained the name of the prescription, the function, the composition, and the name of the prescription book to which the prescription belonged and the year of its publication, final selection of 2231 prescriptions [S1 Table]. Then we standardized the separated and labeled functions of prescriptions into normalized TCM patterns based on TCMeSH, *Dictionary of Traditional Chinese Medicine* and *Terminology specification for common clinical symptoms in Chinese medicine*, farther normalized their definitions.

**Fig 1.**
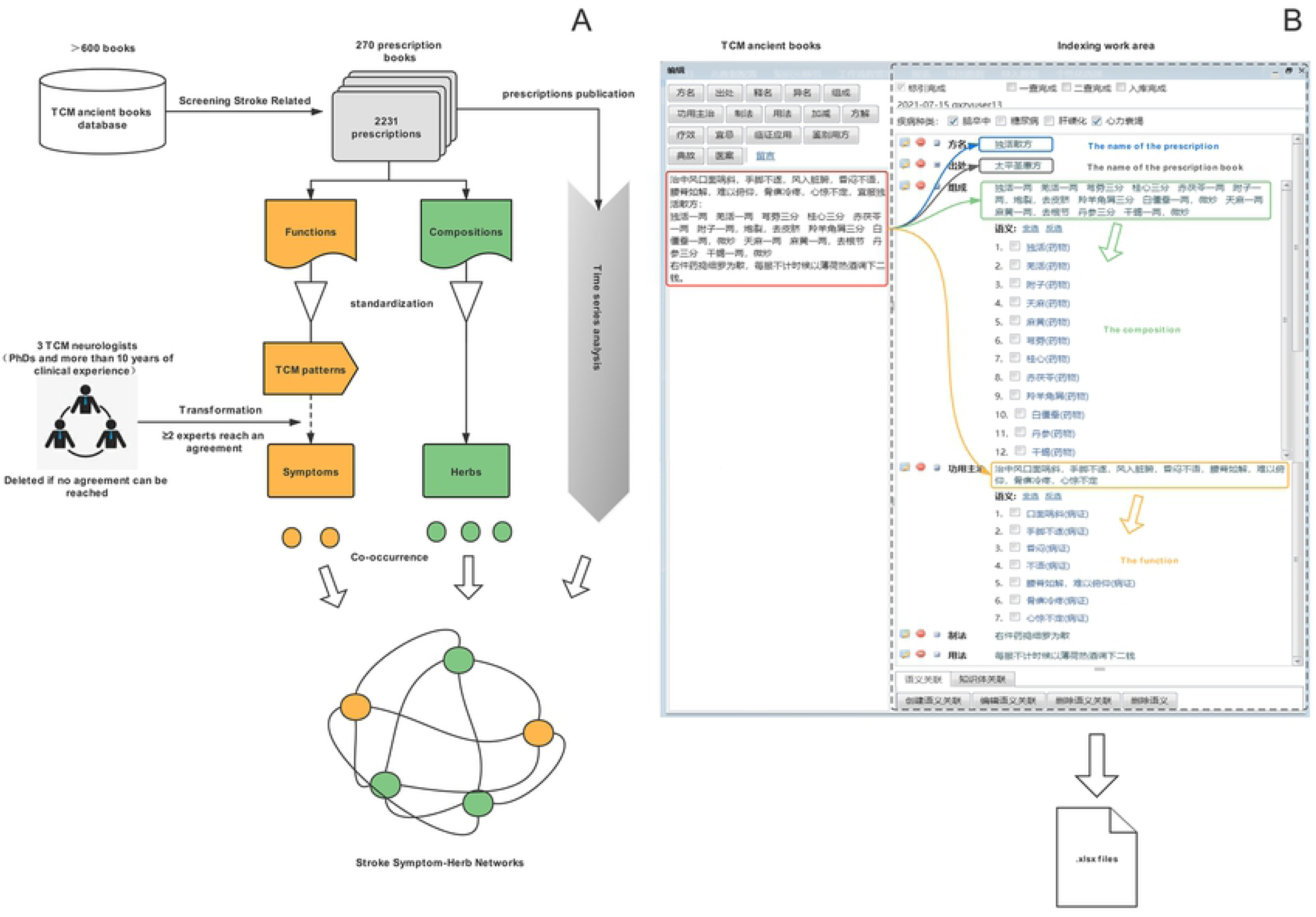
Schematic of the SSHNs construction. (A) The workflow of mining the co-occurrence symptoms and herbs from TCM prescription books for SSHNs. (B) The functions and compositions of the prescriptions are separated and labeled on our TCM ancient books indexing platform.

Subsequently, we referred to the *Neurologic Manifestations* subject terms within MESH and selected 26 symptoms regarding stroke, as well as set 8 symptoms with TCM characteristics. Then, 3 TCM neurologists with PhDs and at least 10 years of clinical experience matched TCM patterns with the 34 selected symptoms, if more than two of them can make an agreement, and the TCM patterns were transformed into symptoms subsequently [S2 Table], with the TCM patterns deleted if no agreement can be reached. Finally, we manually standardized the compositions in the prescriptions into herbs based on national standard *GB/T 31774* which was issued by General Administration of Quality Supervision, Inspection and Quarantine of the People’s Republic of China (AQSIQ) and Standardization Administration of the People’s Republic of China (SAC) in 2015, and some ambiguous herbs that appeared only once and could not be standardized were removed [S3 Table], manual rechecking was conducted to ensure the accuracy of the database.

### Time series analysis

To reflect the evolution of the TCM network by temporal changes, we divided the stroke networks according to the chronology to which the prescriptions belonged. First, we assessed the similarity of frequency curves between the symptoms and herbs using morphological similarity distance (MSD)[4], which was implemented using R language program (R version 3.6.2, RStudio version 1.4.1717), and set to perform similarity calculations at division years of 1, 20, 50, 100, 150, 200, 500, 1000, and 1500. In order to effectively represent the morphological features and trends of the time series, the number of segments needs to be set reasonably. Too few segments will lose critical detail information and morphological features; too many segments will keep more redundant information, increasing the computational complexity.

Finally, we find that, independently of before 600 CE (341-599) as the first chronological period, which may mainly be because of more null values before 600 CE, and every 150 years backward as an interval, the distance no longer changes regardless of the increase in the number of division years, and in Figure 2 the MSD between *Hemiplegia* and all herbs can show that setting the time separation in this way is stable for the whole network, and this formed 11 chronological stages, which were before 600 CE, 600-749 CE, 750-899 CE, 900-1049 CE, 1050-1199 CE, 1200-1349 CE, 1350-1499 CE, 1500-1649 CE, 1650-1799 CE, 1800-1949 CE, and 1950-2000 CE. Also, we were surprised to find that such a separation also has a greater similarity to the historical dynastic separation. Based on the complex network found by the key symptom and herb nodes, the similarity distance of the frequency of occurrence of symptoms and herbs was used for further analysis.

**Fig 2.**
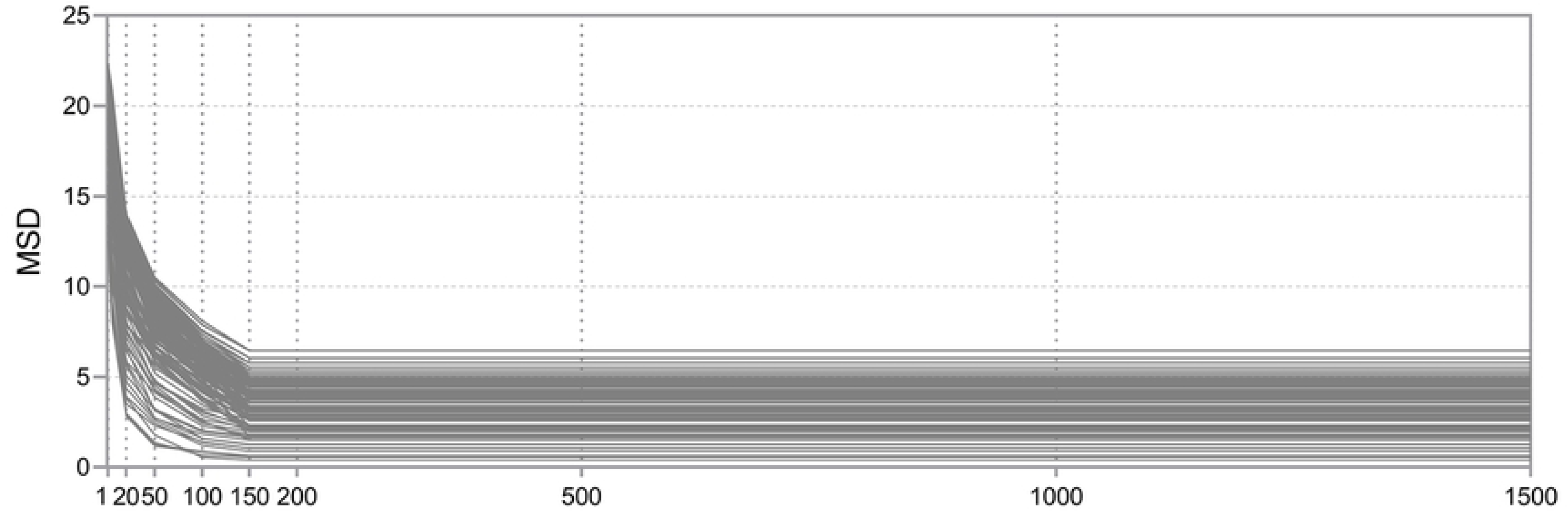
The MSD between Hemiplegia and all herbs at division years of 1, 20, 50, 100, 150, 200, 500, 1000, and 1500.

### Stroke Symptom-Herb Networks construction and analysis

The data were checked by the specialists and crosschecked by the research team for plausibility. After completion of the checkup, all data were prepared (.xlsx files) and subsequently analyzed using the Python (version 3.6) data analysis toolkit “pandas” (version 1.3.0), then the symptoms and herbs were used as nodes, and the edges were connected according to co-occurrence, and the undirected SSHNs was constructed according to each chronological stage, and the Network topological features (including nodes, edges, clustering coefficient, density, degree, the average shortest path length and network centrality, etc.) under each stage were calculated with networkX toolkit (version 2.5.1)[5], and the nodes and edges data were imported into Gephi plots. The genism toolkit (version 4.0.1) was used to calculate the cosine similarity of the symptom sub-networks. We sparse vectors of all herbs by doc2bow work to form a corpus, use TF-IDF model algorithm to process the corpus, calculate the TF-IDF value, get the feature number of token2id, calculate the sparse matrix similarity, build an index, and calculate each group of data of sparse vectors to get the similarity and thus the herb similarity profile of the symptom sub-networks.

## Results and Discussion

### Networks Topology

We connect the nodes of co-occurring symptoms, herbs to edges and build different SSHNs according to different stages [see Figure 3], and the network characteristics recorded in S4 Table reflect the changes of the SSHNs in different stage, which implies the continuous understanding of stroke in Chinese medicine, is it always a gradual progression?

**Fig 3.**
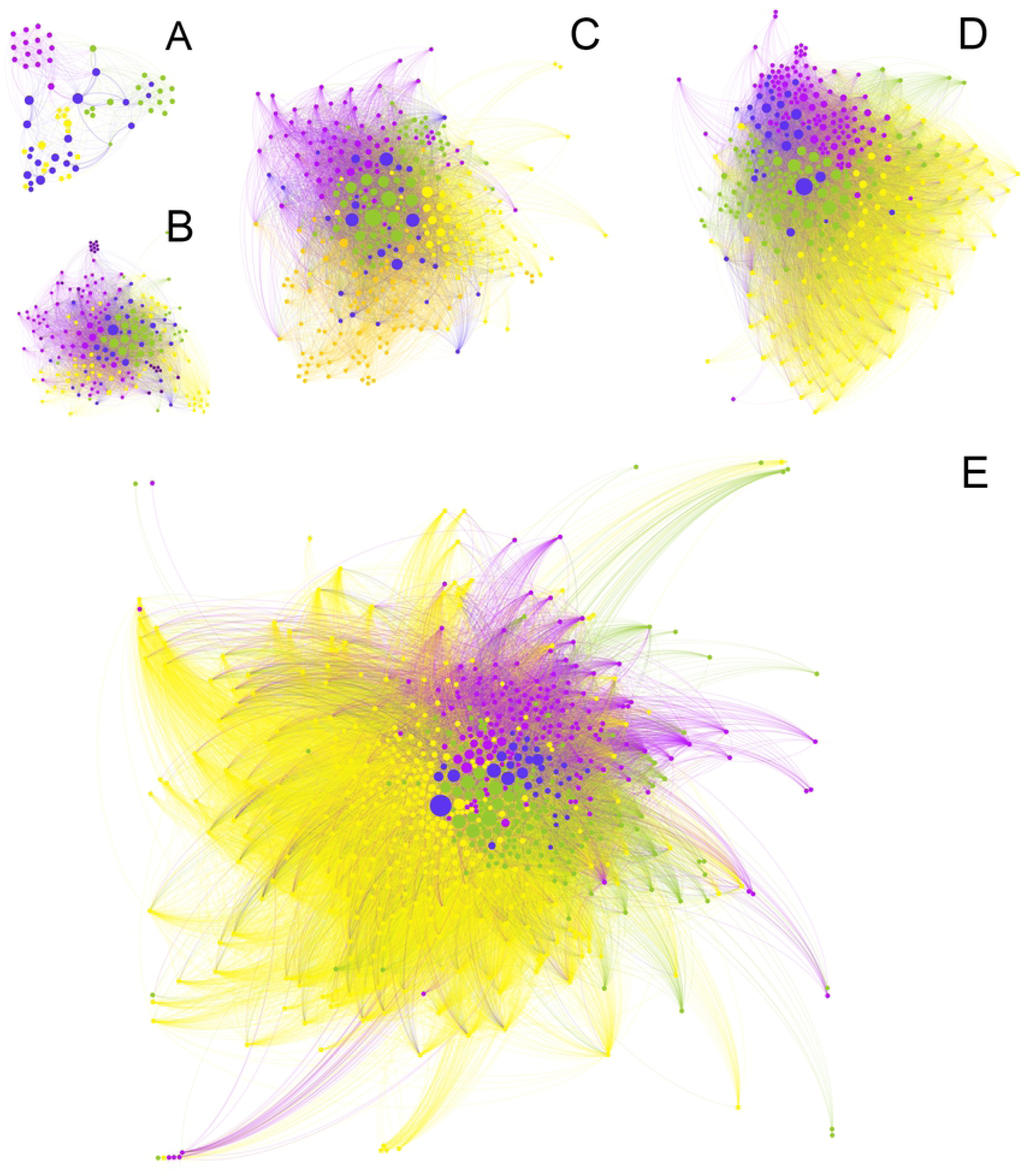
Global SSHNs for different periods. Gephi software was used to connect the symptom and herb nodes in each chronological period based on co-occurrence to gain the network graph under each chronological period where blue nodes are symptom nodes, then herb nodes are yellow, purple, green, etc. The nodes of different colors belong to different communities. The network shows different shape and size at different time periods.

### Nodes and edges

In Figure 4A-B, we can see that the characteristics of nodes and edges indicated they reached their highest number of nodes in 1050-1199 CE, and the highest number of edges in 1800-1949 CE, respectively. Herb node richness peaks and gradually stabilizes over time in 1050-1199 CE, 1200-1349 CE shows a decline in nodes and edges, while from10^th^ century to 12^th^ century shows an upward trend and reaches the first peak in nodes and edges. This period coincides with the Chinese Song Dynasty, which for the first time carried out to separate medical administration and medical education and set up a separate bureau of imperial physicians to take charge of medical education, and the political of medical officials was elevated, and officials collated and proofread antique medical books and published medical writings[6]. Among them, *Taiping Shenghui Fang* and *Shengji Zonglu* were represented by over 16,000 and nearly 20,000 prescriptions[6], while the only official publishing prescription book of the Jin-Yuan period, *Yuyao Yuanfang*, has just over 1,000 prescriptions.

**Fig 4.**
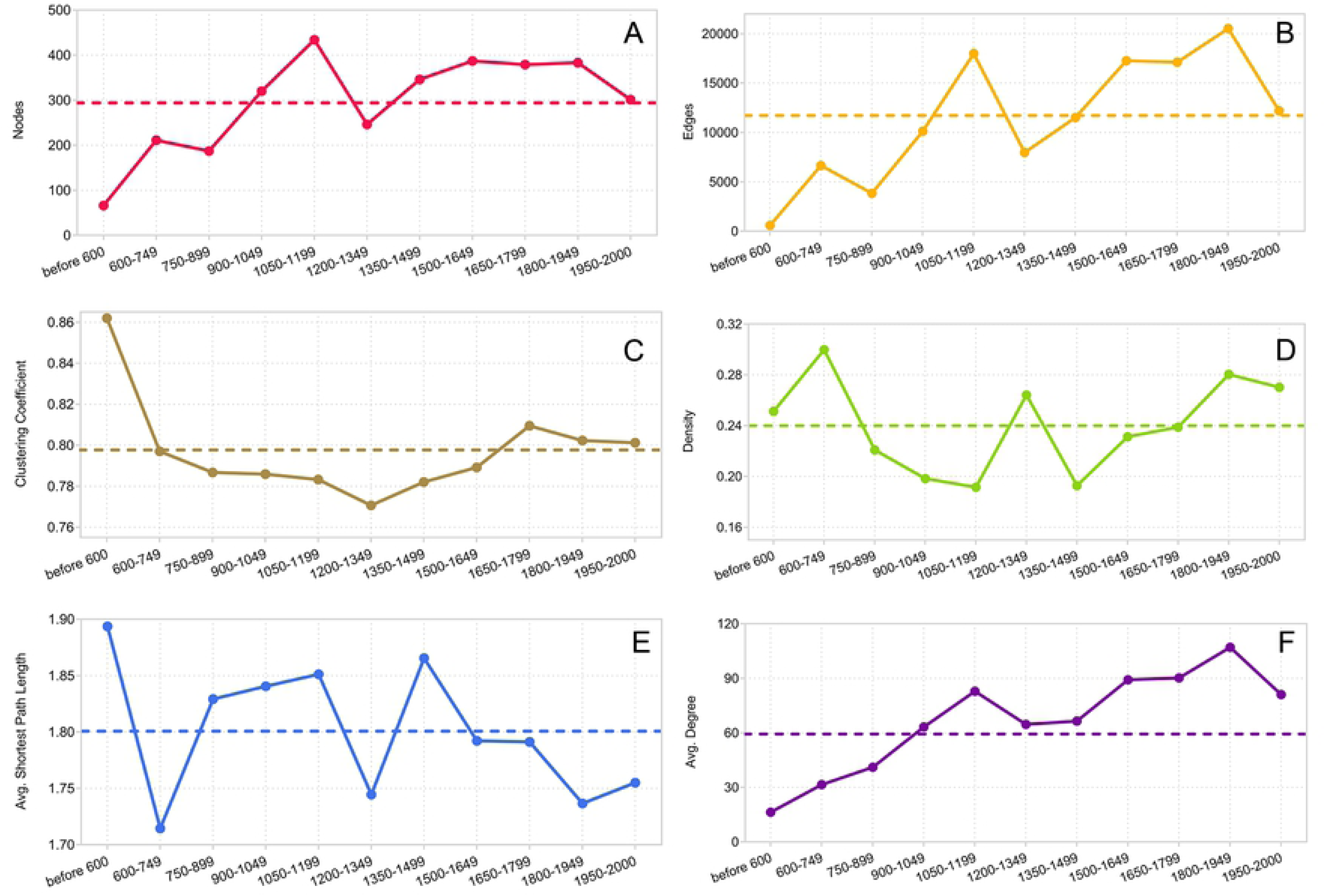
Dynamics of topological parameters for the SSHNs.

However, Chinese folk medicine developed significantly during this period instead, and many Chinese medical practitioners innovated on theories of stroke, laying a key foundation for the second peak that began after 1350 CE. In the 1350-1499 CE, the Ming dynasty officially revised the largest prescription book gathered mainly from folklore in history, *Puji Fang*, in 1406 CE, which contained over 60,000 prescriptions. Until 2000 CE, there was no more large official revision of prescription books, instead folk prescription books developed unprecedentedly, we can see that from 1600 to 1949 CE the nodes remained high-level, showing a smooth peak, at the same time the edges reached a historical peak within this stage of Late Qing Dynasty, showing at which point the SSHNs have best web connections.

### Clustering coefficient and density

Clustering coefficient is a more local measure of cliquishness or how tightly nodes in the graph are connected to each other. The clustering coefficient of the SSHNs always maintains high values (〈C〉> 0.77) in Figure 4C, showing that it is neither a random network nor a tree-like network, and high clustering coefficient phenomenon indicates the SSHNs has a small-world nature. The clustering coefficient decreases gradually from the beginning, coming to its lowest value in 1200-1349 CE, and then gradually increases, falling back after 1650-1799 CE, with a smoother trend at the end. Due to the limited knowledge of stroke in the prior history, despite an enormous increase in nodes in the early to mid-term, edges were not abundant and clustering coefficients continued to decline until after 1200-1349 CE, when it accumulated. This is because the ancient TCM clinicians s had a new understanding of stroke during this stage, creating new theories.

Early TCM clinicians believed the origin of stroke was “external wind”, believing the causes of stroke were related to the external environment, which including the poor living environment and poor protection from cold in ancient times, and this understanding was widely recognized in the Tang and Song dynasties. After entering the Jin-Yuan period, more TCM clinicians found that this understanding was still inadequate and proposed the theory of “internal wind”, believing that the factors of stroke could also originate from internal causes and be closely related to the patient’s own physical characteristics, long-term dietary irregularities, emotional disorders and other risk factors. This interpretation became a widespread consensus among TCM clinicians in the Ming and Qing dynasties after 1350 CE. From the structure parameters of SSHNs for each stage, it can be seen that the network show an intricate topology, which is not the result of randomly generated connection. The network density varies slightly as the SSHNs changes, but always remains above 0.19. Meanwhile, the “external wind” and “internal wind” theories have led to two peaks in Chinese medical theory of stroke[7].

### The average shortest path length

The average shortest path length of the whole SSHNs was 1.801248, as we can see in Figure 4D, showing that any two nodes can be connected by two nodes, and the network diameter is always 3, showing a clear small-world characteristic. In the four stages between the 7^th^ and 12^th^ centuries, the average shortest path length gradually increases and shows a positive correlation with the number of nodes and edges, showing that the network gradually becomes “sparse”, which is closely related to the abundance of herbs in this period. The average shortest path length in the four intervals from 1350 to 1949 CE showed a decreasing trend and a negative correlation with the number of edges, indicating the SSHNs were “aggregated” signaling the network had gradually matured.

### Degree

The number of edges directly connected to the node is the degree, and the average of the degrees of all nodes is called the average degree of the network. In the SSHNs, the trend of the average degree shows an overall upward trend, with a small downward trend during 1200-1499 CE, followed by a small increase. The degree distribution test suggests that the degree distribution does not obey a Poisson distribution. There are a large quantity of nodes with relatively small degrees, and in some stages networks have long-tail characteristics. However, the few nodes with the largest degrees are also not absolutely dominant and do not conform to the power distribution, but some of the larger networks reveal long-tail characteristics, with the whole SSHNs being the most pronounced. Understanding stroke in ancient Chinese medicine was developed, with some new herbs appearing in different chronological stages. Part of herbs continued to be applied in the next stage, therefore, there were some herb nodes with a very low number of occurrences and relatively small degree, showing long-tail characteristics in Figure 5.

**Fig 5.**
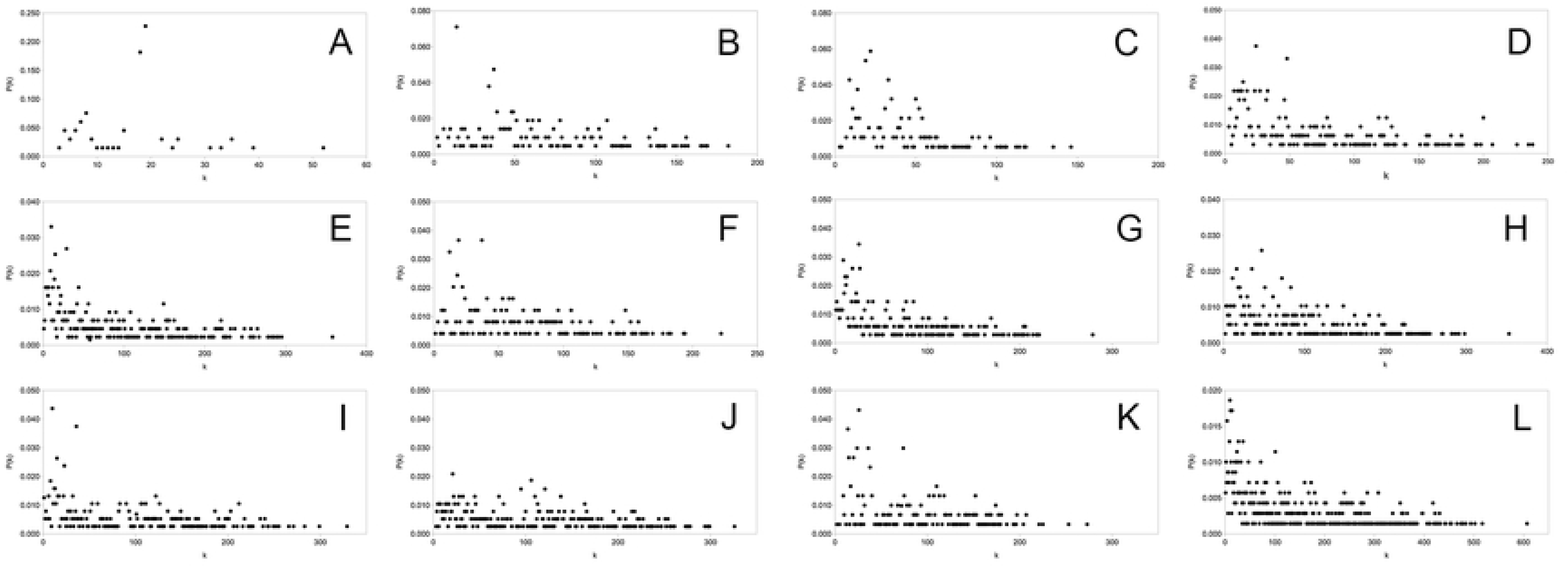
The degree distribution of different periods. Panels A–L: before 600 CE, 600-749 CE, 750-899 CE, 900-1049 CE, 1050-1199 CE, 1200-1349 CE, 1350-1499 CE, 1500-1649 CE, 1650-1799 CE, 1800-1949 CE, 1950-2000 CE, and the whole stages.

In Figure 6 the Upset plot was constructed based on the matrix of herbs in each stage to study the intersection of herbs in different stages [8]. We found the uniquely herbs were added in the 1500-1649 CE with 46, followed by the 1050-1199 CE with 33. These two periods were the Ming and Song dynasties, respectively, which were the two peak periods in the development of TCM stroke theory, and thus unique herbs were added, proving innovative practical exploration of ancient TCM clinicians. Interestingly, there were no new herbs added in the two stages, before 600 CE and 1200-1349 CE, pointing out the herbs in these two stages continued to be used in all later stages, precisely the origin of the two peaks. About intersection, we found that the post-600 CE intersected the most, more than the whole stages, also post-900 CE, showing that the core herb cluster has been initially formed since 600 CE.

**Fig 6.**
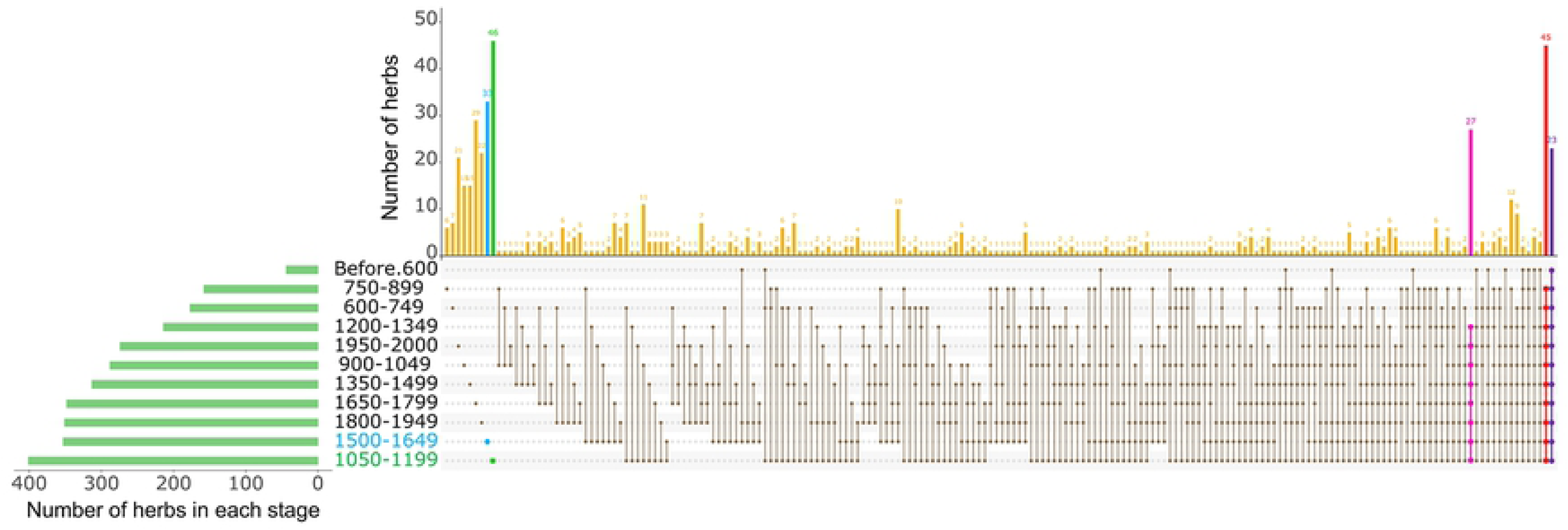
UpSet plot for herbs in all stages. Green strip on the left shows the number of herbs included in each stage. Dots and lines represent subsets of herbs in different stages. The histogram above represents the number of herbs in each subset. Green dot and histogram:1050-1199 CE, blue dot and histogram: 1500-1649 CE, red line and histogram: post-600 CE, pink line and histogram: post-900 CE, purple line and histogram: the whole stages.

### Network centrality

We filtered out critical nodes that always passing through history based on the intersection of symptom and drug nodes across different stages, nodes that were contained in both adjacent stages. It is they that build the major structure of the SSHNs, and that a node in one, or several non-adjacency stages alone will affect its consideration of centrality in the network. Different node centrality metrics, each with its own advantages and characteristics, are available. Therefore, in Figure 7, several centrality methods for the centrality of critical nodes in stroke medication networks were compared by Receiver Operating Characteristic (ROC) curve analysis based on the online website (https://hiplot.com.cn), and we found that degree centrality performed best, followed by betweenness centrality, and finally closeness centrality with eigenvector centrality (S5 Table). This is due to a node that always passing through more stages in the network has more connection, then that node can be assumed to be more central, which leads to the degree centrality becoming the most critical evaluation.

**Fig 7.**
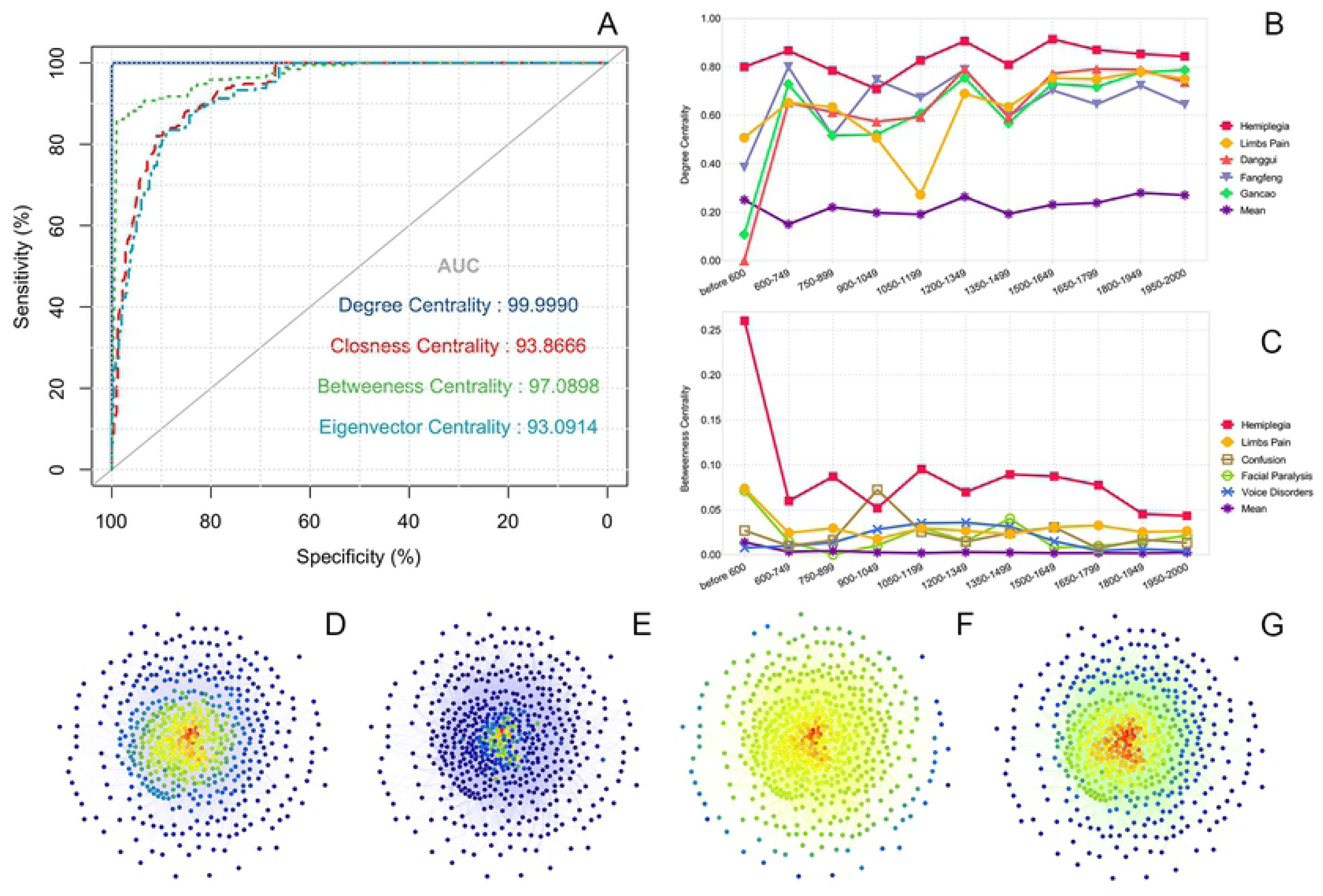
Schematic of Network centrality. (A) Receiver operating characteristic (ROC) curves for 4 centrality methods in comparing the performance of Network centrality. (B) Top five nodes in degree centrality of the whole SSHNs. (C) Top five nodes in degree centrality of the whole SSHNs. (D-G) Gephi is used for the visualization of degree centrality, betweenness centrality, closeness centrality, and eigenvector centrality for the sub-network of *Hemiplegia*.

Degree centrality is the most direct measure of the centrality of a node in network analysis. The larger the degree is, the more important the node in the network [9]. We consider nodes that pass through the whole dynamically SSHNs as core nodes, and they have the highest degree centrality, resulting in the highest AUC values for degree centrality. We find that the top 5 nodes got by ranking nodes based on the three metrics of degree centrality, closeness centrality, and eigenvector centrality are the same: *Hemiplegia*, *Limbs Pain, Danggui*, *Fangfeng*, and *Gancao*. And the top 5 nodes with the highest betweenness centrality are all symptom nodes. The ranking of nodes according to the centrality constantly changes over time, and *Hemiplegia* has always been the most critical node, except for the period from 900 to 1049, during which it was surpassed in betweenness centrality by *Confusion*. Among the herb nodes, *Fangfeng* was the most critical node before 1200 CE, and the centrality of it declined in the later years, while *Danggui* jumped to the top. This is related to the shift from “external wind” to “internal wind” in stroke theory.

Betweenness centrality refers to the proportion of the shortest paths going through a node in all the shortest paths, which show the ability of the node to act as a hub or bridge in the network [10]. In the SSHNs, the vast majority of nodes have a lower betweenness centrality and the value of some nodes are zero and some of the symptom nodes have higher betweenness centrality. Top 5 nodes with the highest betweenness centrality include *Hemiplegia*, *Limbs Pain*, *Confusion*, *Facial Paralysis* and *Voice Disorders,* playing a key pivotal role in the SSHNs. *Hemiplegia* is the most important node in almost all periods of the SSHNs, indicating that *Hemiplegia* is the most common symptom of stroke and that TCM has the deepest understanding and rich experience in treating it. TCM attaches great importance to *Limbs Pain*, especially in stroke, and lots of herbs have been proved in pharmacological experiments to have pain-relieving effects. *Hemiplegia*, *Confusion*, *Facial Paralysis* and *Voice Disorders* are 4 of the 5 items in the prehospital stroke scale, Conveniently-Grasped Field Assessment Stroke Triage (CG-FAST).

In 2021, the top neurology journal Lancet Neurol published an article titled *Comparison of eight prehospital stroke scales to detect intracranial large-vessel occlusion in suspected stroke (PRESTO): a prospective observational study* [11], and found that the CG-FAST performed best in the 8 prehospital assessment scales tested in this study. However, it is regrettable that another item *Oculomotor Paralysis* on the scale does not show higher centrality in the SSHNs, which is related to the lack of attention to this symptom in ancient TCM.

Degree centrality only uses the local features of the network, but a node with a high number of connections does not mean that it is at the core of the network. Closeness centrality determines the centrality of a node by calculating the reciprocal of the sum of the shortest paths to all other nodes; a node whose closeness centrality is high is close to many nodes, thus showing that it is of more significant importance [9]. As well as betweenness centrality, closeness centrality makes use of features of the entire network, where a node is in the overall structure. If the shortest distance from a node to any other node in the graph is small, then its closeness centrality is high. Closeness centrality is closer to geometric centrality than betweenness centrality.

Eigenvector centrality means that the importance of a node depends both on the number of its neighboring nodes (nodes degree) and the importance of its neighboring nodes [12]. Taking the sub-network of *Hemiplegia* as an example, it can be seen in Figure 7D-G that a node may have many neighboring nodes, but it’s eigenvector centrality may not be necessarily high, because the nodes it connects may have low eigenvector centrality. Conversely, a node with high eigenvector centrality does not mean that it has many neighbors either, just that they are all critical nodes. Eigenvector centrality has the lowest AUC because it differs the most from the way degree centrality is determined.

### Sub-networks

We extract the corresponding sub-network according to different symptoms, which comprise the herbs used for treating that symptom interconnected as nodes. The purpose of comparing and analyzing the topological differences and similarities between different sub-networks is to analyze the rules of medication use in the treatment of different symptoms in TCM stroke.

For these subnetworks, we recorded values for the number of nodes and edges, clustering coefficient, network density, network diameter, shortest path, mean degree, and network centrality (see S6 Table). The larger sub-networks are *Hemiplegia*, *Limbs pain*, *Somatosensory Disorders*, *Sialorrhea*, *Headache*, *Muscle Cramp*, *Facial Paralysis*, *Voice Disorders*, *Confusion*, *Poor Appetite*, and *Emotion Disorders,* and they have more number of nodes, edges, and average degree. The sub-networks of *Olfaction Disorders and Taste Disorders* are smaller, while their network density and clustering coefficients are higher. The clustering coefficient of all sub-networks is greater than 0.74, with the means very close to the entire SSHNs. The average shortest path and average density values for each sub-network are lower than the whole network. We also found that most of the symptom nodes with significant weight or high centrality correspond to larger sub-networks, and interestingly, Neither the weight nor the centrality of node *Poor Appetite* in the SSHNs is prominent, while the number of nodes and edges, average degree, shortest distance, and network density of the *Poor Appetite’s* herb sub-network are all ranked higher. This shows that TCM attaches great importance to the symptom *Poor Appetite*, including loss of appetite, anorexia, inability to eat, and so on. The experience of TCM in the treatment of these symptoms is also more abundant, with *Renshen*, *Baizhu*, *Fuling* and *Gancao* all having high degrees, and they form a prominent TCM prescription for dietary disorders, SiJunZi Tang.

By calculating the centrality of nodes in sub-network and the edge betweenness, we can know core herbs and the critical couplet herbs for the different symptoms [S7 Table]. Every sub-network shows different structural and dynamic evolutionary features, and in Figure 8 we can see the characteristics of the two sub-networks of *Hemiplegia* and *Confusion*. *Hemiplegia* is the most central node in the SSHNs, with significantly higher weights, degrees, and centrality than the other nodes, and therefore the sub-network for *Hemiplegia* and the global network must show similar dynamics in their temporal evolution. There are two distinct periods of increase in the scale of the sub-network for *Hemiplegia*, 900-1199 CE and 1350-1649 CE, then the number of nodes stabilized at high levels thereafter. The number of edges rises again significantly between 1800 and 1949, which is congruity with the SSHNs. In contrast, the scale of the sub-network for *Confusion* increased significantly in 900-1199 CE, and then remained stable at lower levels after a significant decline in 1200-1349 CE. Before 600 CE, the herbs for *Confusion* do not form a network, but several groups of scattered nodes, which cannot form a network, and the value of clustering coefficient is zero. However, with the progressive understanding of *Confusion* in TCM, the average clustering coefficient and the average shortest path of the sub-network for *Confusion* are both higher than the sub-network for *Hemiplegia* after 750-899 CE.

**Fig 8.**
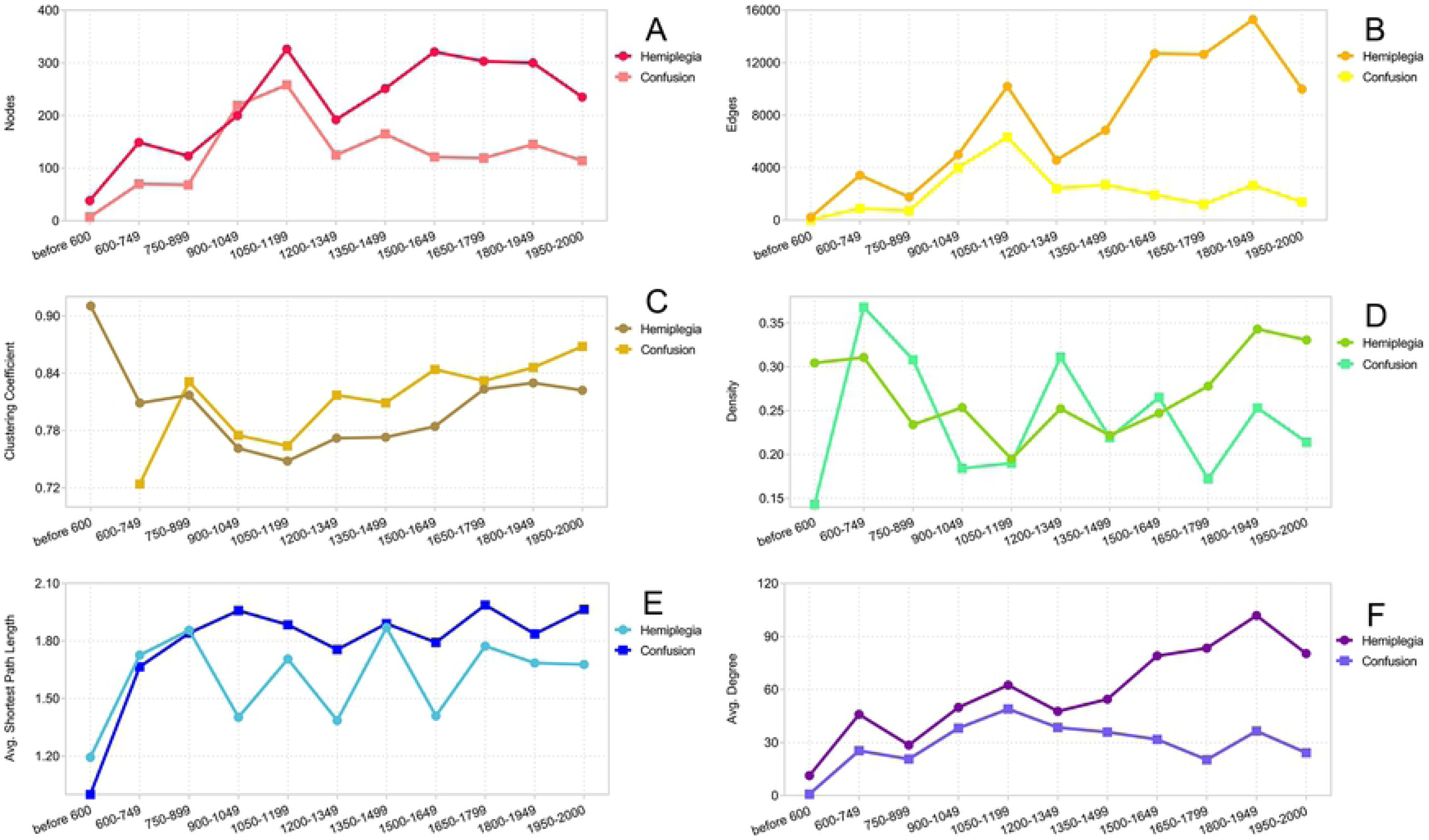
Dynamics of topological parameters for the two sub-networks of Hemiplegia and Confusion.

### Sub-network similarity

To further explore the similarity between the sub-networks, we calculated the cosine similarity of the sub-networks of different symptoms via the gensim toolkit in python 3.6 [S8 Table], and then their spearman correlations were calculated by using R language, and heatmap was drawn through the heatmap package as shown in Figure 9 [13].

**Fig 9.**
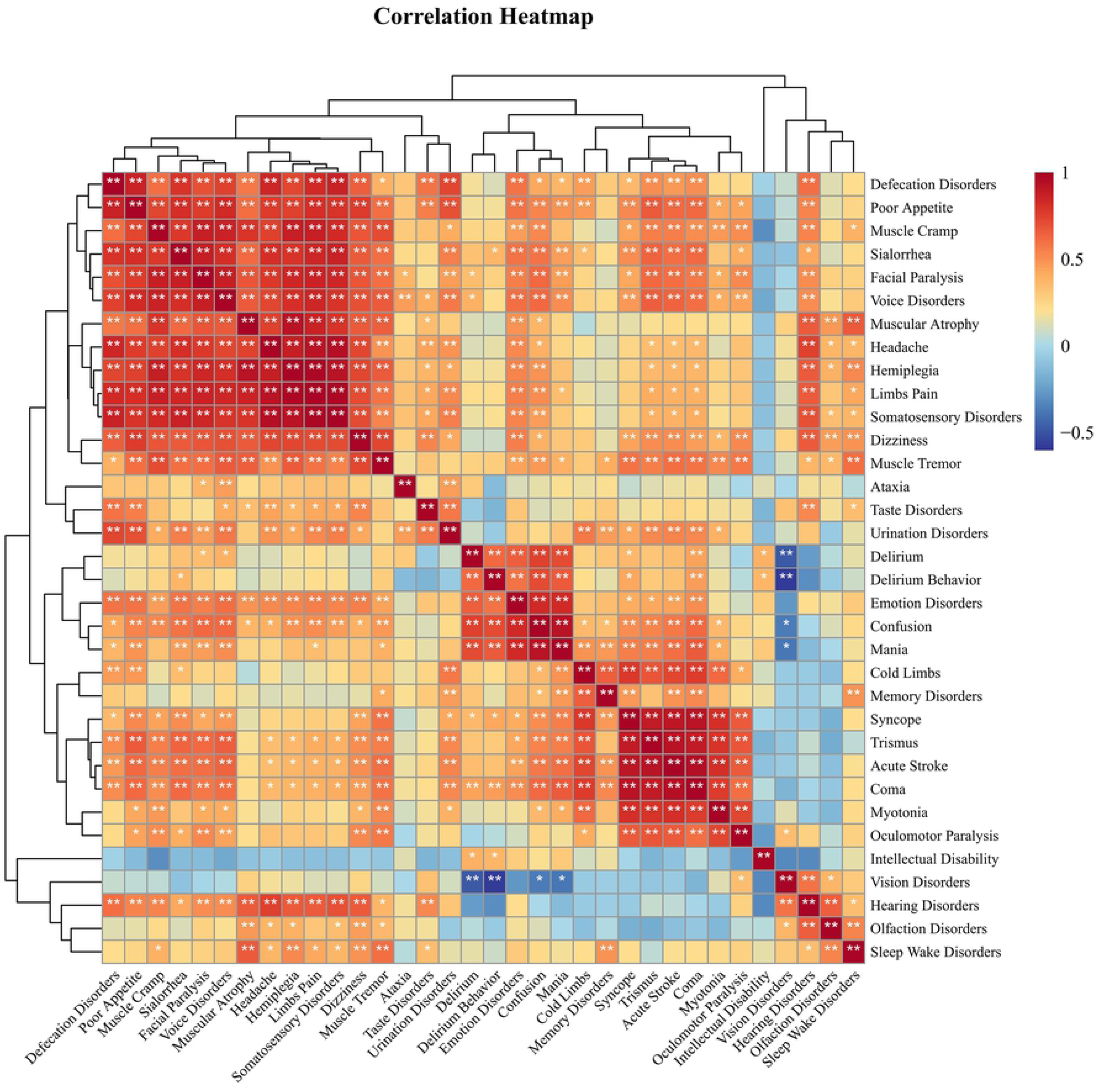
A heatmap of the cosine similarity of the herb sub-networks of different symptoms was calculated using the Spearman correlation coefficient.

There is a significant positive correlation between the sub-network of *Hemiplegia*, *Limbs Pain*, *Somatosensory Disorders* and *Headache*. The correlations between the sub-network of *Coma*, *Syncope*, *Trismus* and *Acute Stroke* are also very significantly positively. That is because the herbs for symptoms are highly correlated, and meanwhile *Acute Stroke* is often accompanied by *Syncope* or *Coma*. While the sub-network of *Coma*, *Syncope* and *Acute Stroke* shows negative correlations with the sub-network of *Hearing*, *Olfaction*, and *Vision Disorders*, this is due mainly to the fact that the former symptoms are mostly in a *Coma* state, while the latter are mostly manifested under clear consciousness.

Likewise, the sub-network of *Delirium* and *Delirium Behavior* negatively correlated with the sub-network of *Hearing*, *Olfaction* and *Vision Disorders*, especially *Vision Disorders*. Whereas studies showed that *Vision Disorders* is a well-recognized and important risk factor for *Delirium* after stroke[14]-[15]. The reason for this situation is that the herbs with tranquilizing spirit by heavy settling, such as *Zhushafen*, *Xijiao*, etc., are widely used in treatment of *Delirium* and other manifestations of mental abnormality. While Chinese herbal medicine for the treatment of *Hearing*, *Olfaction* and *Vision Disorders* has high efficacy of warming meridians and unblocking collaterals, such as *Chuanwu* and *Fuzi*. The category of those herbs has strong opposite medicinal properties, cold-property against warm-property, resulting in an enormous difference in the sub-network.

### Community Detection

In a randomly connected network, the pattern of connections between nodes should be identical and therefore should not show systematic local density differences. This means that there will be no community. Studies show that networks in the real world all have obvious community structures. After comparing two community partition results based on modularity and label propagation algorithm, we choose the Louvain non-overlapping community detection based on modularity optimization method built in Gephi. Modularity, as a metric that can measure the quality of community division, was proposed by Newman and Girvan in 2004[16]. Automatic calculation processing and optimization analysis of modularity were used to divide the community structure of the SSHNs according to each chronological period. Resolution is a parameter used to control the number of communities divided. The smaller the resolution set, the more communities are. In this study, when the resolution is set to 0.9-1.1, the modularity of each stage reaches the highest value, which is 0.426, 0.139, 0.241, 0.156, 0.144, 0.137, 0.164, 0.155, 0.182, 0.156, 0.179, respectively, so that the community division reach its optimum [S9 Table]. Basically, the networks of different stages are all divided into 3-5 communities. Subsequently, we analyzed the dynamic community evolution, and some nodes that only appeared at a single stage and then disappeared were eliminated. By observing the nodes in different communities of different stages, we find that the community where *Hemiplegia* and its associated nodes, such as *Limbs Pain*, *Somatosensory disorders*, *Niuxi*, *Shinan*, *Yiyiren,* are located form a relatively permanent historical trend, the upper strip in Figure 10. In addition, *Sialorrhea* also forms a relatively permanent combination with its related nodes, that is the strip shown at the bottom in Figure 10. Among them, some symptom nodes such as *Syncope* and *Coma*, and some herb nodes, such as *Niuhuang*, *Shexiang*, *Shuiyin* and *Bingpian* have been closely related to *Sialorrhea* all the time.

**Fig 10.**
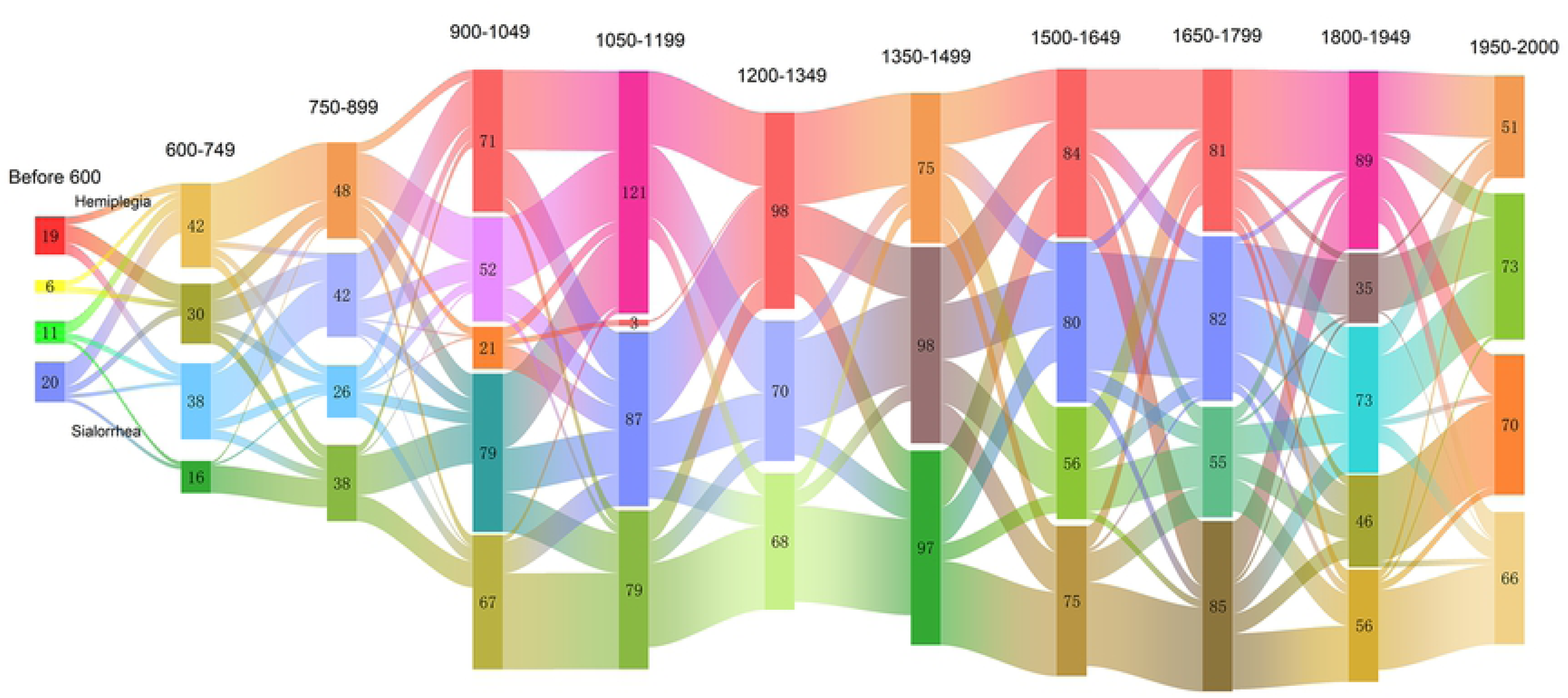
Sankey diagram for the evolution of divided communities based on modularity in different stages.

### Time series analysis

To further uncover the deeper relationships between symptoms and medications in the SSHNs on the timeline. MSD with certain morphological recognition ability is chosen in this paper to analyze and calculate the similarity between two time series of symptoms and herbs. The similarity of time series is measured by their distance, and the smaller the distance between them, the more similar the two sequences are; oppositely, the larger the distance, the less similar. Euclidean distance is one of the most widely used tools for similarity calculation, and MSD is a method for similarity calculation based on the classical Euclidean distance, considering the specific distribution factors of the difference of each dimension of the object to be calculated. The existing research results show that the MSD can evaluate the similarity by combining the size and shape of the trajectories [17].

Considering vectors as objects, the morphology similarity distance is proposed to estimate similarity. The morphology similarity distance between two *n*-dimensionality vectors ***L****_i_* = ( *l_i_*_1,_ *l_i_*_2_, …, *l_i_*_n_ ) and ***L****_j_* = ( *l_j1_*, *l_j2_*, …, *l_j_*_n_ ) is defined as:

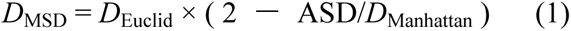

where *D*_Euclid_ is the Euclid distance, *D*_Manhattan_ is the Manhattan distance, ASD is the absolute sum of the differences as follows:

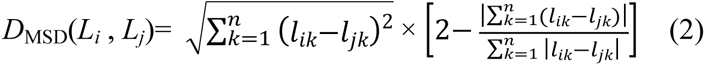

Based on the performance of symptom and drug nodes in the SSHNs, from which we screened 22 symptoms and 52 herbs, these were the nodes with the highest frequency, the highest node degree, namely the most significant degree centrality and the nodes that were consistently present for the SSHNs. We measured the MSD of the frequency curves between symptom and herb (S10 Table), and found that the ranking of time-series similarity for *Hemiplegia* and *Voice Disorders* was more similar to its degree centrality ranking, with *Chuanxiong* and *Danggui*, both of which have high edge betweenness and are also usually in a same network community, commonly playing a critical role in activating blood and resolving stasis. As we can see in Figure 11A, their MSD curves are very similar. The results of 45 prescriptions containing *Chuanxiong* and *Danggui* analyzed by Gumbel copula function model showing that the *Chuanxiong* and *Danggui* have a strong nonlinear correlation in stroke, and the correlation is positive [18]. Studies have shown that their extracts exert neuronal protective effects by promoting endogenous proliferation of neuroblastoma cells and production of neural differentiation factors after ischemic injury from stroke in rats [19]- [20].

**Fig 11.**
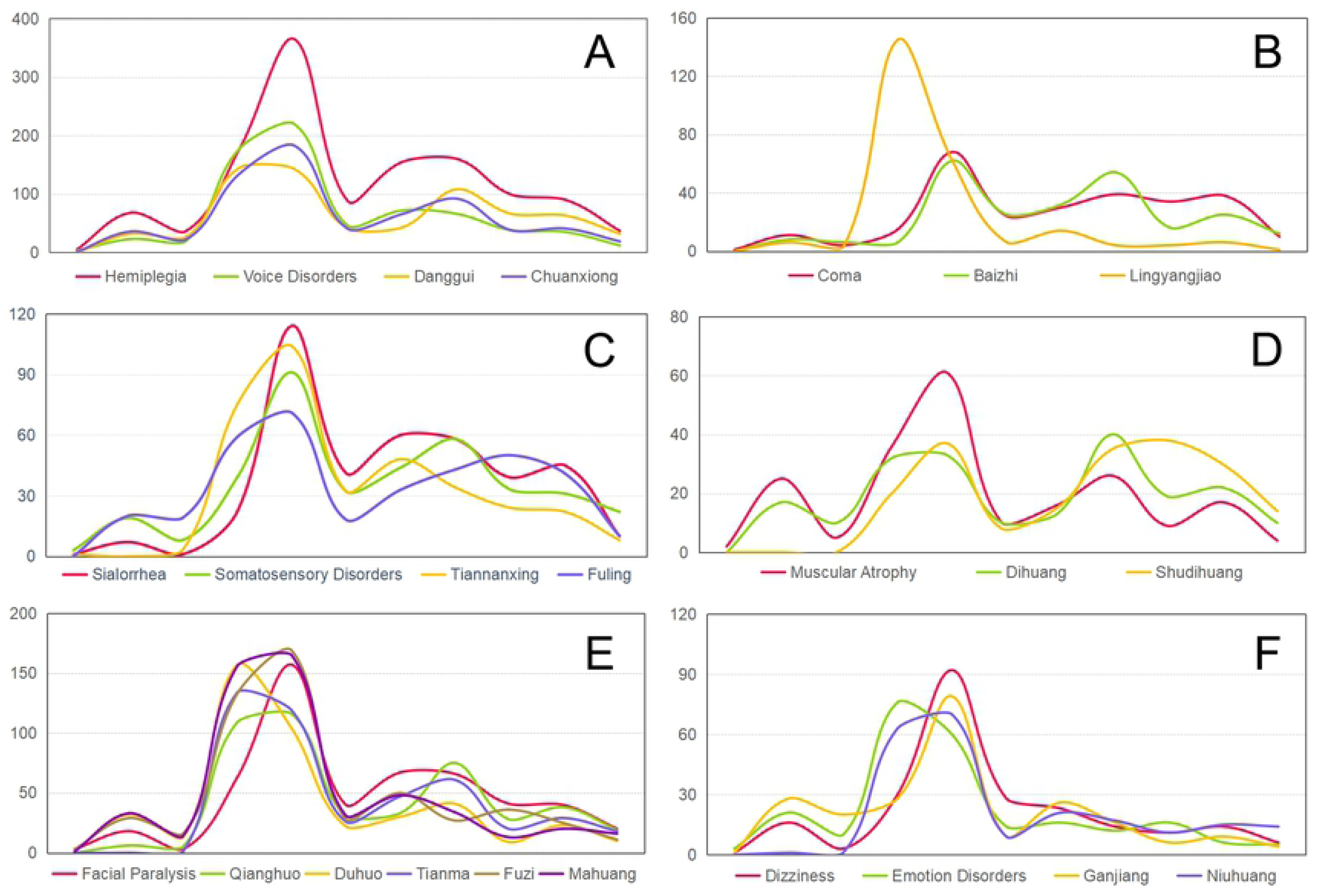
MSD curves for the time-series similarity of symptom and herb frequencies were measured. (A) *Hemiplegia*, *Voice Disorders*, *Chuanxiong* and *Danggui*. (B) *Somatosensory Disorders*, *Sialorrhea*, *Fuling* and *Tiannanxing*. (C) *Dizziness*, *Emotion Disorders*, *Ganjiang* and *Niuhuang*. (D) *Facial Paralysis*, *Qianghuo*, *Duguo*, *Tianma*, *Fuzi* and *Mahuang*. (E) *Coma*, *Baizhi*, and *Lingyangjiao*. (F) *Muscular Atrophy*, *Dihuang*, and *Shudihuang*.

As we can see in Figure 11B, in the ranking of herbs with the highest similarity of *Somatosensory Disorders* and *Sialorrhea*, *Fuling* and *Tiannanxing* were both significantly elevated. In TCM, phlegm is the sign of *Sialorrhea* obstructing meridians and collaterals is the key pathogenic mechanism, and the main composition of Ditan Tang with *Fuling* and *Tiannanxing* is an extremely important TCM prescription for treating phlegm blocking the meridians in stroke disease.

The top ten ranking of the herbs with the highest similarity in *Facial Paralysis* and *Coma* differs completely from the ranking of degree centrality. Among them, *Qianghuo*, *Duguo*, *Tianma*, *Fuzi* and *Mahuang* have the smallest MSDs with *Facial Paralysis*, and their frequency curves are always close to each other [Figure 11C]. That is because they all have the function of dispersing and dissipating pathogenic external wind for *Facial Paralysis* in the “external wind” theory of TCM stroke. Besides, *Baizhi* and *Lingyangjiao* have the effect of opening orifices and unblocking block to treat *Coma*.

We were surprised to find that the top ranking of herbs with the highest similarity in *Dizziness*, *Emotion Disorders* were similar, especially *Ganjiang* and *Niuhuang* [Figure 11E]. In the previous study it can be found that *Dizziness* is not very similar to the sub-network of *Emotion Disorders*, and in the total network *Ganjiang* and *Niuhuang* have a very low edge betweenness, and never belong to the same community, showing that they are poorly correlated, but they are both closely associated with *Dizziness* and *Emotion Disorders. Ganjiang* and *Niuhuang* are two herbs with sizeable differences, both with opposite medicinal properties, and the combination of these two herbs will provide us with fresh ideas than before.

In Figure 11F one can see that *Dihuang* and *Shudihuang* are very eminent for *Muscular Atrophy*, especially *Dihuang*, which shape is very similar to *Muscular Atrophy* in the latter part of the time axis. This also shows a significant correlation with the second peak in stroke, the “internal wind” theory of TCM. A systematic evaluation of the efficacy of Catalpol in experimental acute focal ischemic stroke in 25 studies of 805 animals showed that Catalpol, one of the effective components of *Dihuang*, has a neuroprotective effect on experimental acute focal ischemic stroke, mainly by reducing oxidative response, inhibiting apoptosis, inhibiting inflammatory response and autophagy[21].

## Conclusion

Chinese medicine attaches great importance to the symptoms of diseases, especially in stroke, and ancient TCM clinicians have a rich knowledge of stroke symptoms and have accumulated a lot of valuable experience in prescriptions and herbs. In this study, we deeply explored the symptom-herb relationship of stroke on the timeline by constructing the SSHNs and enriched our knowledge related to stroke symptom-herbs, to employ novel technical means to uncover the deep information of stroke prescriptions in the history of TCM, and trying to establish associations with the more standardized MESH terms, for the next step of herb network pharmacology and active ingredient experimental.

## Acknowledgments

We gratefully acknowledge Dr. Jingling Chang, Lingbo Kong, Xinke Yang for the time they spent matching TCM patterns with symptoms.

## Supporting Information

**S1 Table. The original database of the SSHNs containing 2231 prescriptions.** First column contains ID, second column is the Chinese name of the prescriptions, third column is the function of the prescriptions, fourth column is the composition of the prescriptions, fifth column is the Chinese name of the prescription books, sixth column is the year of publication of the books.

**S2 Table. Transformation from TCM pattern terms to symptom terms.** First column contains various prescription functions in the original prescription books, second column contains TCM pattern terms got by standardizing the content of the first column, third column is the definitions of TCM pattern terms in Chinese, fourth column shows the TCM pattern terms that have been translated into English, fifth column shows the definitions of TCM pattern that have been translated into English, sixth column is the reference source of these definitions, the seventh column is the 34 symptom terms selected by TCM neurologist experts, Y in the eighth column means the symptom term refer from MESH, ninth column is the definitions of these terms in MESH.

**S3 Table. Data for all herbs.** First column contains various herb names in the original prescription books, second column contains TCM herbs standardized by *GB/T 31774*, fourth, fifth and sixth columns are the standardized herb in Pinyin, English and Latin respectively.

**S4 Table. Dynamics of topological parameters for the SSHNs.** First column contains the years, second column is the number of nodes, third column is the number of edges, fourth column is the clustering coefficient, fifth column is the networks’ density, sixth column is the shortest path length, seventh column is the average degree, eighth column is the networks’ diameter, ninth column is the average degree centrality, tenth column is the average closeness centrality, eleventh column is the average betweenness centrality.

**S5 Table. ROC for performance comparison of four centrality measurement methods.**

**S6 Table. The topological parameters of the symptom sub-networks.** First column contains 34 symptom terms, the second column is the number of nodes, third column is the number of edges, fourth column is the clustering coefficient, the fifth column is the networks’ density, sixth column is the shortest path length, seventh column is the average degree, eighth column is the networks’ diameter, ninth column is the average degree centrality, tenth column is the average closeness centrality, eleventh column is the average betweenness centrality.

**S7 Table. Centrality of all symptom sub-networks.**

**S8 Table. The cosine similarity between different symptom sub-networks.**

**S9 Table. The resolution set in the community detection and its corresponding modularity and the number of communities.**

**S10 Table. The MSD of the frequency curves between symptoms and herbs.**

